# Sampling biases obscure the early diversification of the largest living vertebrate group

**DOI:** 10.1101/2022.05.09.491144

**Authors:** Struan Henderson, Emma M. Dunne, Sam Giles

**Affiliations:** School of Geography Earth and Environmental Sciences, University of Birmingham, Birmingham B15 2TT, UK; Department of Earth Sciences, Natural History Museum, Cromwell Road, London SW7 5BD, UK

**Keywords:** diversity, sampling standardisation, Actinopterygii, fossil record bias, Palaeozoic

## Abstract

Extant ray-finned fishes (Actinopterygii) dominate marine and freshwater environments, yet their spatiotemporal diversity dynamics following their origin in the Palaeozoic are poorly understood. Previous studies investigate face-value patterns of richness, with only qualitative assessment of potential biases acting on the Palaeozoic actinopterygian fossil record. Here, we investigate palaeogeographic trends and apply richness estimation techniques to a recently-assembled occurrence database for Palaeozoic ray-finned fishes. We reconstruct patterns of local richness of Palaeozoic actinopterygians, alongside sampling standardised estimates of ‘global’ diversity. We identify substantial fossil record biases, such as geographic bias in the sampling of actinopterygian occurrences centred around Europe and North America. Similarly, estimates of diversity are skewed by extreme unevenness in the abundance distributions of occurrences, reflecting past taxonomic practices and historical biases in sampling. Increasing sampling of poorly represented regions and expanding sampling beyond the literature to include museum collection data will be critical in obtaining accurate estimates of Palaeozoic actinopterygian diversity. In conjunction, applying diversity estimation techniques to well-sampled regional subsets of the ‘global’ dataset may identify accurate local diversity trends.

## Introduction

There are around 32,000 species of living ray-finned fishes (actinopterygians), amounting to over half of extant vertebrate diversity, and split roughly evenly between marine and freshwater environments (1). Ray-finned fishes originated in the Palaeozoic, which saw major evolutionary events and changes in the vertebrate fauna, such as the emergence of jaws (2), the rise of actinopterygians (3), and the move onto land (4). Despite these pivotal changes, and a long history of research on actinopterygians, there are relatively few macroevolutionary studies investigating diversity trends in their early evolution, and all examine face-value patterns of taxonomic richness (3,5–9).

Few studies investigate the suitability of the Palaeozoic ray-fin record for investigating diversity patterns, or potential biases. Notably, biases may impact the marine and freshwater record differently – late Palaeozoic Lagerstätten influence freshwater osteichthyan diversity more than marine (5). Low taxonomic diversity in the Devonian followed by an explosive increase in the early Carboniferous is generally interpreted as representing a genuine biological signal (3,10). Some authors qualitatively suggest that low Permian diversity is linked to the rarity of suitable deposits (11), or attribute the decline in richness among freshwater taxa to the loss of extensive Euramerican freshwater habitats (5). Other authors propose that the consistent ecomorphologies in typical Palaeozoic actinopterygians hint at constraints on diversification into new ecologies and habitats and thus low richness (10). To date, previous studies only present face-value counts of actinopterygians through time without employing recent advances in methodologies to estimate diversity trends. An exception to this (5) performed coverage-based rarefaction to compare the Permian and Triassic as a whole, rather than to estimate diversity trends through time.

Assessing the degree to which fossil record biases affect interpretations of richness is critical to obtaining an accurate estimate of diversity trends (12–14). These biases can be geological (15,16), geographic (17–19), or anthropogenic (20,21) in nature, and recent analyses show that ‘global’ fossil records are intimately linked to the spatial extent of that record (17,18). Various statistical methods attempt to tease apart bias from genuine changes (e.g. classical rarefaction and residual modelling), though not without complications (e.g. classical rarefaction can flatten diversity patterns (22–25). Recent years have seen the application of Shareholder Quorum Subsampling (SQS), also termed coverage-based rarefaction (25,26), to palaeobiological occurrence databases (17–19,27–31) as a means of deducing trends in palaeodiversity through time. As SQS subsamples intervals to equal levels of completeness it returns more accurate relative richness estimates between sampled intervals than size-based rarefaction (23), although is still susceptible to some biases (21,24). Principally, SQS estimates can have a significant evenness signal (21,24,32), which may be particularly important for datasets that are biased in ways that skew the evenness of frequency distributions within sampled intervals. A new richness estimator, squares (33), estimates higher richness when there are numerous rare taxa (i.e. singletons) and when common taxa are especially abundant. Squares is more robust to uneven distributions than SQS, though falls short when the ratio of richness counts to total number of taxa within intervals is very high (24).

Until recently, no comprehensive through-Palaeozoic occurrence database existed (9), with previously-published databases limited in scope or not updated (3,5). Here, we apply coverage-based sampling standardisation to a newly-assembled occurrence database of Palaeozoic actinopterygians to examine patterns of diversity, the suitability of the dataset, and the likely extent and impact of sampling biases.

## Methods

### Data preparation

Global occurrences of Palaeozoic Actinopterygii (9) were screened for taxonomically indeterminate occurrences and scale- and teeth-based occurrences. After removal, this resulted in a dataset of 1,611 occurrences of 468 species (belonging to 225 genera), from 507 unique geographic localities. All occurrences were assigned to intervals of roughly equal length (~9 Ma), determined by either combining shorter intervals (e.g. Kasimovian [3.3 Ma] and Gzhelian [4.8 Ma] = Kasimovian and Gzhelian [8.1 Ma]), or splitting longer intervals (e.g. Visean [15.8 Ma] = early Visean [Chadian-Holkerian; 8.7 Ma] and late Visean [Asbian-Brigantian; 7.1 Ma]; boundary based on the age of the Dunsapie basalt, see (34)). The cleaned dataset was used for local richness and diversity estimation. All analyses were conducted within R 4.1.0 (35).

### Alpha diversity (local richness)

Species per locality were counted as a measure of alpha diversity (local richness (36)). Modern coordinates for these localities were translated into palaeocoordinates using the R ‘chronosphere’ package (37). Local richness was then subset by marine and freshwater environment (brackish environments were included in marine counts) and plotted against palaeolatitude. Additionally, palaeogeographic maps showing local richness were produced in ‘chronosphere’ (37) for each interval. It is uncertain whether some Permian localities (Pastos Bons – Brazil; Deep Red Run, Dundee, McCann Quarry, Pond Creek, South Dakota State Cement Plant Quarry – USA; Sobernheim – Germany) are Artinskian or Kungurian in age, and these localities are therefore plotted in both palaeogeographic maps.

### Sampling standardisation and diversity estimation

Coverage-based sampling standardisation (22,26,38,39) was used to estimate global diversity patterns via the R package iNEXT (version 2.0.19 (40)), following the procedure outlined in Dunne *et al.* (28). The data were rarefied by geographic locality by analysing incidence-frequency matrices of the occurrence data. Extrapolated estimates were limited to no more than twice the observed sample size (40). Coverage-standardised richness was computed at genus level using roughly equal length bins, at quorum levels 0.3, 0.4, 0.5 for genus-level analysis and up to 0.6 quorum for species-level analysis; higher quorums were unattainable. Devonian bins were excluded due to the very small sample-sizes and low levels of coverage. Rank abundance distributions and size- and coverage-based rarefaction curves were generated for each interval to investigate the reliability of coverage-based rarefaction estimates.

Squares extrapolated estimates of genus and species richness were conducted in R by applying Alroy’s equation (33), following the same procedure as Allen *et al.* (30).

## Results

### Alpha diversity (local richness)

Local richness is generally low in the Devonian (figure 1), with only one locality containing more than three genera (Paddy’s Valley, Gogo Formation, Frasnian, Australia). Levels of local richness are highest in the Carboniferous, particularly around the Serpukhovian-Moscovian boundary (figure 1), before declining steadily in the latest Carboniferous (Kasimovian and Gzhelian) and early Permian (Cisuralian). Notable localities contributing to the mid-Carboniferous peak include Glencartholm (Scotland, late Visean, marine), Ardenrigg (Scotland, Bashkirian, freshwater), Longton (England, Bashkirian, marine) and the Bear Gulch localities (USA, Serpukhovian, marine) (figure 1a). Sampling of marine and brackish palaeoenvironments in the latest Carboniferous (Kasimovian and Gzhelian) and earliest Permian (Asselian and Sakmarian) is very poor. Freshwater localities are also poorly sampled in the Artinskian and Kungurian, yielding very low richness, while richness and sampling also remains low in marine deposits. In the latest Permian (Wuchiapingian and Changhsingian), marine localities generally have much higher genus counts than freshwater localities.

**Figure 1.**
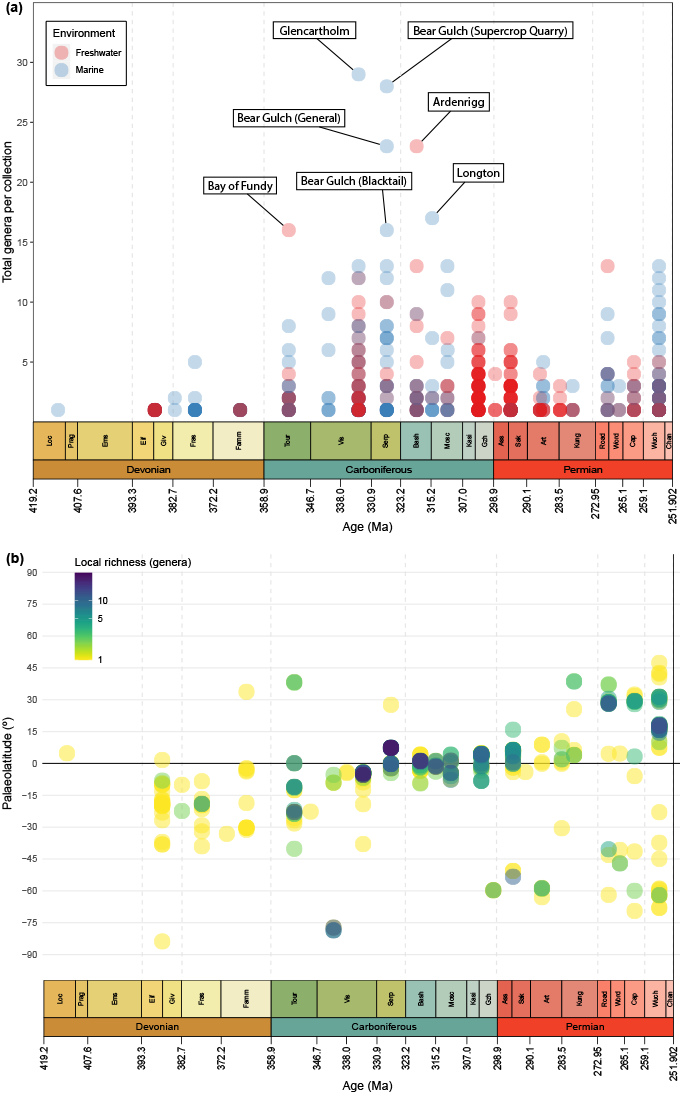
Local richness (number of species per fossil locality) of actinopterygians through the Palaeozoic. (a) Local richness plotted by environment, separated by freshwater (red) and marine (blue; incorporates brackish occurrences). Colour saturation (transparency) indicates density of localities, and the most genus-rich localities are labelled. Note that purple indicates contemporary localities with similar diversity of marine and freshwater actinopterygians. (b) Palaeolatitude of localities through time, with local richness indicated by colour (yellow localities have low richness, progressing through green to the most diverse localities in indigo).

### Palaeomaps and geographic spread

#### Devonian

Despite their earliest occurrence being just north of the palaeoequator (*Meemannia*, Lochkovian, South China), actinopterygians are known almost exclusively from southern palaeolatitudes in the Devonian (figure 1b; figure 2a). Only two other northern hemisphere occurrences are reported (*Cheirolepis*, Givetian, Svalbard (41); *Krasnoyarichthys*, Famennian, Russia (42)). The majority of taxa occur at low palaeolatitudes (0° to −30°), with a small number just crossing into the mid-palaeolatitudinal band (−30° to −60°). A clear outlier, near the southern palaeopole (−83.81°), is the recently-reported *Austelliscus ferox* from Brazil (43).

**Figure 2.**
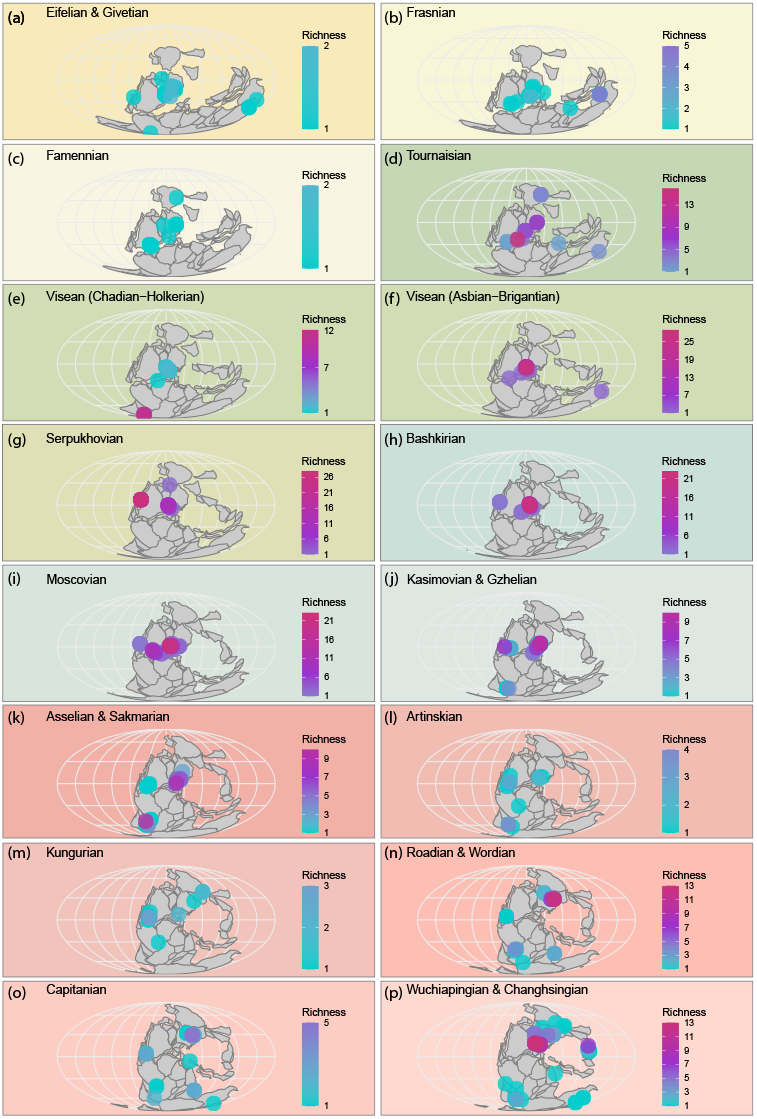
Palaeomaps showing palaeogeographic spread and local richness of individual localities through time plotted in roughly even-length intervals. Colour scales correspond to the richness of localities, ranging from low (light blue) to high (pink) richness.

Devonian actinopterygian occurrences mirror both continental configurations—the majority of landmasses and shallow seas were palaeoequatorial and in the southern hemisphere (44)—and the broader Devonian fossil record (18,45,46). Givetian and Eifelian occurrences are dominated by European (especially Scottish) deposits, with limited contributions from the USA, Australia, the Antarctic and Brazil (figure 2a). In contrast, Frasnian occurrences (figure 2b) are dominated by the Australian Gogo Formation, with fewer occurrences from Europe and North America, and a single occurrence from Iran. The USA dominates Famennian occurrences (figure 2c), with additional occurrences from Russia, Greenland and Belgium.

#### Carboniferous

In general, Carboniferous localities have both higher local richness and a broader palaeolatitudinal spread than in the Devonian, although are generally still restricted to low and southern palaeolatitudes (figure 1b). Most Tournaisian localities are clustered at low palaeolatitudes around the southern edge of Euramerica in regions that correspond to present-day Canada, USA, UK, and European Russia. Localities with lower local richness are found in Australia, Turkey, and Siberia. In contrast, early Visean (Chadian-Holkerian) low-to mid-palaeolatitudes are depauperate (figure 2e), although the Waaipoort Formation in South Africa (−78°) represents the richest high-palaeolatitude locality of the entire Palaeozoic.

For much of the rest of the Carboniferous, local richness greatly increases while palaeolatitudinal spread decreases. Other than single occurrences from Australia and the USA, all late Visean actinopterygians are clustered in the UK and Ireland, including the highly diverse Glencartholm locality (figures 1b and 2f). Similarly, in the Serpukhovian (figure 2g), only a single occurrence is found outside a 20° palaeolatitudinal band centred around the palaeoequator encompassing UK localities, a single Belgian locality, and the speciose Bear Gulch localities. Geographic spread continues to decline in the Bashkirian (figure 2h) and Moscovian (figure 2i), with all occurrences within 10° of latitude of the palaeoequator. Again, localities only are only known in Europe (Belgium, Czechia, France, Ireland, UK) and North America (Canada, USA). The only latest Carboniferous (Kasimovian and Gzhelian; figure 2j) locality outside of this band is the −60° Gzhelian Ganigobis Shale, which outcrops in South Africa and Namibia, albeit with low local richness. Broadly, Carboniferous actinopterygian palaeolatitudinal distribution matched other contemporaneous groups (18,46).

#### Permian

Compared to the Carboniferous and Devonian, Permian occurrences generally display a broader geographic spread (reflecting increases in the broader fossil record (18,46)) but lower local richness. The extent of palaeogeographic sampling in the Asselian and Sakmarian (figure 2k) is greater than the Kasimovian and Gzhelian, with more occurrences at higher palaeolatitudes, including the diverse Uruguayan fauna from Rio Negro (−53°). The Artinskian (figure 2l) is the most depauperate interval of the Palaeozoic outside of the Devonian, despite a comparatively high palaeogeographic spread: the locality with the highest local richness, Loeriesfontein, contains only four genera. Contrary to most other Palaeozoic intervals, there are very few European Artinskian localities.

From the Kungurian (figure 2m) onwards, localities occur across the broadest palaeolatitudinal spread of the entire Palaeozoic. This includes the first sampling of northern mid-palaeolatitudes since the Tournaisian. Wordian and Roadian localities (figure 2n) with the highest local richness are found in Russia, centred around 30° palaeolatitude, although less diverse occurrences are seen at high southern palaeolatitudes in Brazil, India, and Zimbabwe. In contrast to most other intervals, only two depauperate localities occur near the palaeoequator. This trend continues into the Capitanian (figure 2o), where localities yielding few genera are found across a wide range of palaeolatitudes, with very few at equatorial latitudes, and most diversity stems from Russia.

The Wuchiapingian and Changhsingian interval (figure 2p) has the broadest geographic spread in sampling of the Palaeozoic, possibly due to intensive research focus on the Permo-Triassic mass extinction event (47,48). Numerous localities are spread from southern mid-to high-palaeolatitudes, including opposing sides of the palaeopole (present-day South Africa and Australia). Notably, this interval contains the first Palaeozoic actinopterygians from the eastern Palaeotethys (present-day China) aside from a single Lochkovian occurrence. Northern low-to mid-palaeolatitudes have the highest local richness, stemming from assemblages in the UK and Germany, Russia, and Greenland.

### Palaeodiversity estimates

#### Coverage-based rarefaction

Estimates of relative genus richness using coverage-based rarefaction (figure 3a) suggest a gradual overall decline in diversity through the Carboniferous, with a sharp rise then subsequent fall in the Permian. Richness levels decrease from the Tournaisian through to the late Visean (the most intensely sampled interval of the Carboniferous), before peaking in the Serpukhovian. The remainder of the Carboniferous is marked by a decline, with the lowest observed values in the Kasimovian and Gzhelian, another intensely sampled interval. Richness estimates rise slightly across the Carboniferous-Permian boundary, followed by a precipitous rise in the Kungurian, where the highest levels in the Palaeozoic are reached. A steady decline marks the remainder of the Permian.

**Figure 3.**
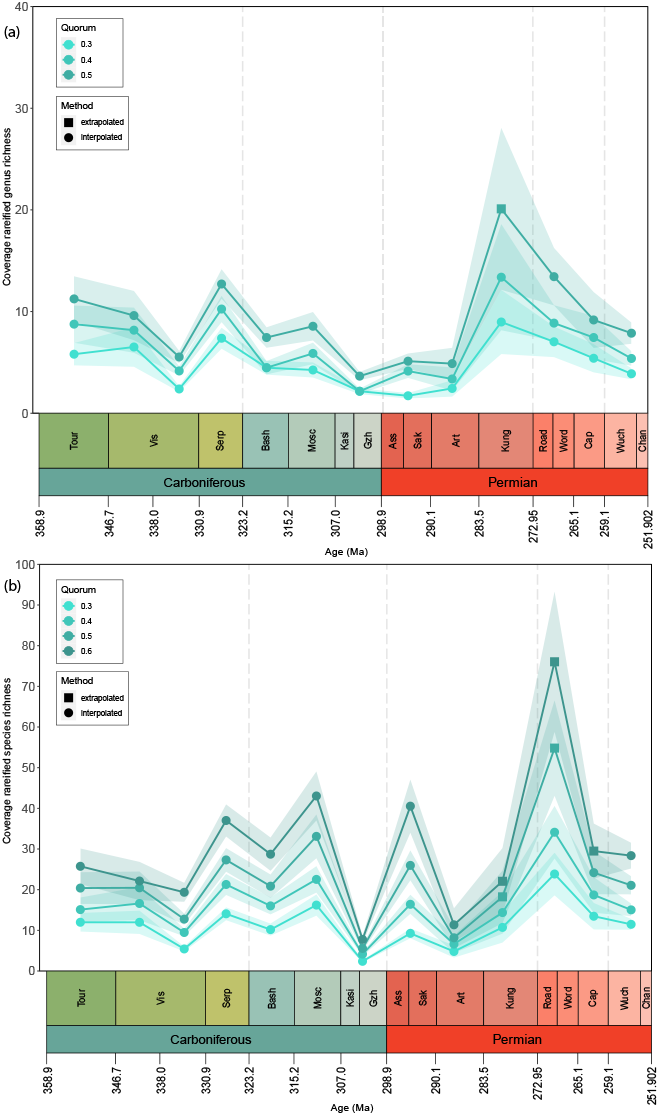
Coverage-based rarefaction estimates of Carboniferous and Permian actinopterygian diversity at (a) genus- and (b) species-level, showing estimates for different quorum levels in different colours from low quorums (0.3) in light blue to higher quorums (0.6) in darker blue. The shaded areas for each quorum are confidence intervals of estimates. Devonian intervals removed (see Methods). Estimates were either interpolated (circles) or extrapolated (squares) up to twice the reference sample size (40).

Coverage-rarefied estimates of species richness differ notably from genus estimates (figure 3b). Overall, estimates of species diversity generally increase, albeit irregularly, across the Carboniferous until a crash in the Kasimovian and Gzhelian, followed by two distinct peaks and declines in the Permian. Species richness initially decreases through the Tournaisian and Visean and increases in the Serpukhovian, with a drop into the Bashkirian and subsequent rise into the Moscovian, which represents the highest richness levels of the Carboniferous. This peak is immediately followed by a Kasimovian and Gzhelian trough. Levels rise steeply in the Asselian and Sakmarian followed by another abrupt drop in the Artinskian. There is only a modest rise into the Kungurian, with the major peak in species-level estimates seen in the Roadian and Wordian. A relative decrease in the Capitanian is followed by a minor decline through the Wuchiapingian and Changhsingian.

#### Squares

Squares diversity estimates contrast starkly with coverage-based rarefaction estimates: where coverage-based rarefaction returns low estimates, squares estimates are generally high. Squares-extrapolated genus richness estimates (figure 4a) gradually increase throughout the Devonian and into the Tournaisian. Early Visean estimates drop back to Famennian levels, before gradually rising in the late Visean to Serpukhovian. A slight decrease into the Bashkirian is followed by a steeper decline in the Moscovian. The highest estimates thus far are seen in the latest Carboniferous with a further increase into the Asselian and Sakmarian, followed by a precipitous drop in the Artinskian. Richness estimates rise in the Kungurian and marginally in the Roadian and Wordian before dropping in the Capitanian. The latest Permian (Wuchiapingian and Changhsingian) is the most diverse interval of the Palaeozoic.

**Figure 4.**
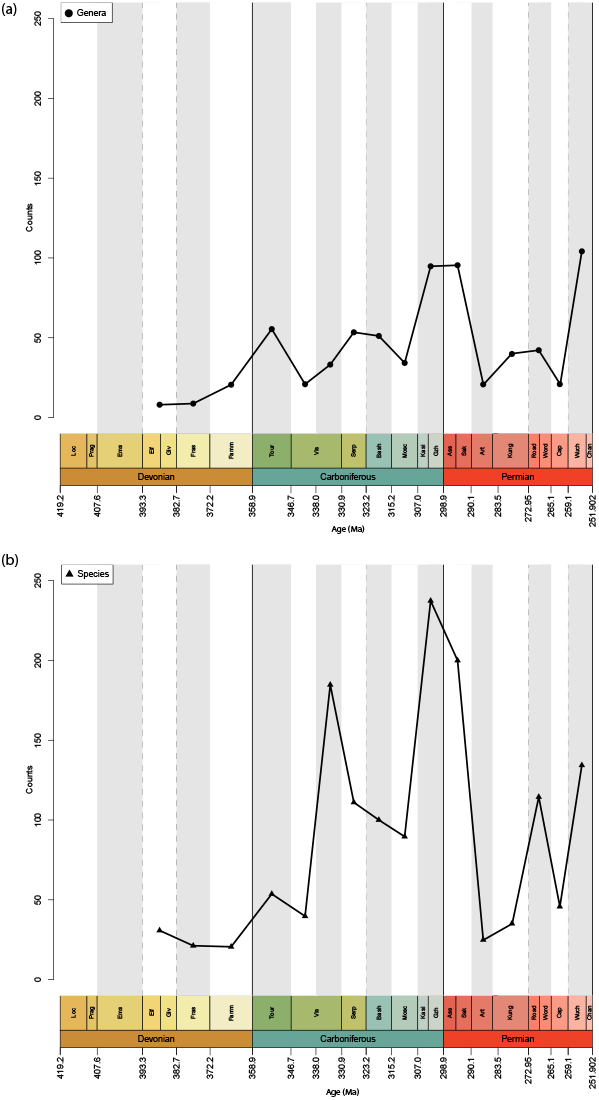
Squares diversity estimates of Devonian to Permian (a) genus (circles) and (b) species (triangles) richness, plotted at the midpoints of equal-length intervals.

Squares-extrapolated species richness trends (figure 4b) differ again from genus richness estimates. The pattern in the Devonian is in direct contrast, with estimates decreasing from the mid-through to late-Devonian in the Givetian and Eifelian, although a rise is observed across the Devonian-Carboniferous. As with genus estimates, the early Visean has lower richness than the Tournaisian. Species richness estimates rise significantly in the late Visean, quadrupling relative to the early Visean. The Serpukhovian sees a moderate decrease in richness, and this trend continues throughout the Bashkirian and Moscovian. Richness rises sharply in the Kasimovian and Gzhelian to the highest level of the entire Palaeozoic. A slight decrease across the Carboniferous-Permian boundary is followed by a precipitous drop in the Artinskian. Richness increases slightly in the Kungurian, recovers further in the Roadian and Wordian, declines again in the Capitanian, and finally increases in the latest Permian.

## Discussion

### Biogeographic trends and biases

Palaeozoic actinopterygian occurrences are overwhelmingly geographically biased towards the northern hemisphere: fewer than 9% of known localities (52/586) are from the southern hemisphere (9). Certain regions are notably underrepresented throughout the Palaeozoic, such as the northern, eastern and southern Palaeotethys (present-day Middle East, south and east Asia, north Africa) and the northern coastline of Laurussia (present-day Siberia, Kazakhstan and interior of Asia). Sampling through much of the Devonian and Carboniferous is limited to a narrow band around the palaeoequator, largely corresponding to present-day Europe and North America (figure 2), which also contain localities with the highest local richness (9). The most diverse localities trend from low-to mid-palaeolatitudes through the Palaeozoic, essentially tracking the migration of North America and Europe (figure 1b). Reporting new taxa from underrepresented regions (41,43) will have major implications for palaeogeographical spread, patterns of diversity, and interpretations of ray-finned fish evolution, especially in the face of taxonomic revisions invalidating many existing generic referrals (49,50).

Ideally, rarefaction curves for sampled intervals should be close to asymptote before performing diversity estimation techniques to ensure that future sampling will not drastically alter face-value counts of richness. Inspection of the Palaeozoic ray-finned fish record suggests this condition has not yet been reached (figure S1). In the short term, increased sampling of the most under sampled intervals will improve comparability. However, research focus on taxa from well-sampled regions that are languishing undescribed in museum collections (51,52) is also vital for attaining accurate estimates of actinopterygian diversity in the Palaeozoic, particularly at local scales.

Both marine and freshwater deposits are recorded throughout the Palaeozoic, with the number of sampled marine and freshwater deposits roughly tracking each other through much of the Carboniferous. However, marine palaeoenvironments are scarce in the later Palaeozoic. This long-recognised Permian imbalance (10,11,53) also extends back into the late Carboniferous (figure 1a). The near-complete lack of marine deposits suggests that low marine diversity in this period is linked to a geological bias and relative absence of these rocks rather than a true biological signal. There is certainly a change in the sampling of terrestrial vertebrates from aquatic to dryland terrestrial environments across the Carboniferous-Permian (54), and a similar change may explain the drop in sampling of Permian actinopterygians. Concurrent with this environmental shift is a noticeable palaeogeographical expansion: rather than being restricted to palaeoequatorial regions, Permian occurrences are reported from much higher and lower palaeolatitudes. It is unclear to what extent this represents a shift in sampling regime rather than an ecological expansion.

### Palaeozoic actinopterygian diversity patterns

Changes in local richness largely track changes in ‘global’ (gamma) raw diversity (9), with the exception of the latest Carboniferous and earliest Permian (figure 1). In the late Carboniferous and early Permian, high levels of sampling (localities and equal-area grid cells (9)) of isolated localities with low alpha diversity drives high ‘global’ diversity, with few contributions from diverse assemblages. These richness patterns are drastically different to those reported for Palaeozoic tetrapods (28), and the overall decrease from the Carboniferous to Permian contrasts the biodiversification of invertebrates over the same period (55).

In contrast to coverage-rarefied diversity estimates, extrapolated estimates from squares analysis return very similar trends to face-value counts of richness (3,5,9). These differences persist regardless of whether sampling is via equal length intervals or geological stages and are likely due to taxonomic biases (see below). This recalls recent work on Palaeozoic tetrapods, which found that diversity patterns among reptiles and synapsids changed significantly depending on the quorum levels or use of squares (56). For example, coverage-rarefied actinopterygian diversity decreases from the Tournaisian to the late Visean in contrast with previous hypotheses (3,6,9), yet both the face-value counts and squares estimates increase significantly from the early to late Visean. There is consensus however, in the high diversity of the Serpukhovian (3,9), indicating genuine diversity, though the vast majority of this is driven by the highly-diverse Bear Gulch fauna.

Trends into the Pennsylvanian also differ, with the greatest difference in the diametrically opposed estimates for the Kasimovian and Gzhelian, which is attributable to the ways in which the methods estimate diversity. The same is also true for the Asselian and Sakmarian and late Permian. Coverage-rarefied diversity estimates depend on the attainable level of coverage, and examination of abundance distributions (figure S2) and rarefaction curves (figure S3) reveals that at higher coverage, the Kasimovian and Gzhelian would most likely represent one of the most diverse intervals. Squares, however, estimates higher richness when there are many singletons and when common taxa are especially common (24), and these intervals fulfil both of these criteria. The combined presence of superabundant taxa and numerous singletons results in these conflicting estimations.

Taxonomy also plays a key role. The observed rise in early Permian species-level diversity estimates in both analyses and face value readings (9) reflects the presence of numerous species of few genera (namely *Amblypterus* and *Paramblypterus*). The problem of superabundant genera is not unique to actinopterygians; such genera are known to bias other osteichthyan groups (57). In contrast, Kungurian estimates are based on very few occurrences of monospecific genera, and sampling of a high number of genera at low quorums results in high—yet unreliable—genus-level coverage-rarefied diversity estimates. The extremely high Roadian and Wordian species-level estimates in both analyses, not reflected at genus-level, can also be explained by high numbers of singletons and relative absence of common genera.

### Unevenness in the actinopterygian fossil record

Coverage-based rarefaction techniques produce the most reliable richness estimates when rank abundance does not differ considerably between samples, even when samples have comparable face-value richness (22–26,32,38). Unevenness in abundance distributions can therefore heavily influence the reliability of diversity estimates. Rank abundance distribution plots for Palaeozoic actinopterygian genera and species indicate extreme unevenness within intervals and variation in evenness between intervals (figure S2). Some intervals (e.g. Kasimovian and Gzhelian) contain one or two taxa with more than 60 occurrences, a handful with between 30 and ten occurrences, and a long tail of singletons or doubletons; others (e.g. Frasnian) have a more even distribution. Differences can even arise between the genus- and species-level abundance distributions in the same interval: in the Asselian and Sakmarian most species-level diversity stems from multiple species of two genera (*Amblypterus* and *Paramblypterus*), resulting in low genus estimates at lower quorums, but higher species-level estimates due to the more even abundance distributions (compare figures S2a and S2b; S3f and S3i).

Much of this imbalance is driven by ‘waste-basket’ genera erected by monographic descriptions (58–61), despite a wide range of varied morphologies and extensive temporal and geographic ranges within genera (9,62,63). These ‘waste-baskets’ serve to concentrate frequency counts of the most common genera, contributing to unevenness in the abundance distribution and distortion of coverage-based rarefaction estimates (22,23). The intervals most heavily biased towards superabundant taxa are the late Visean (*Elonichthys*: 54/266 occurrences; *Rhadinichthys*: 54/266 occurrences), Kasimovian and Gzhelian (*Elonichthys*: 65/230; *Sphaerolepis*: 60/230), Asselian and Sakmarian (*Paramblypterus*: 53/154 occurrences; *Amblypterus*: 30/154 occurrences), and Wuchiapingian and Changhsingian (*Palaeoniscum*: 66/225 occurrences; *Platysomus*: 26/225 occurrences). As coverage-based rarefaction produces lower estimates when evenness is low (23), highly uneven intervals have low richness estimates at lower quorum levels (figure 3; figure S3). In contrast, at high quorums, where more taxa in the abundance distribution can be sampled, uneven intervals receive much higher richness estimates (see exponential rise in the rarefaction curves of uneven intervals at high coverage; figure S3).

‘Waste-basket’ taxa may also mask true diversity: the dominance of highly abundant taxa means that a high proportion of sampled taxa consists of these few taxa, likely contributing to lower diversity estimates. Revisionary taxonomic work, such as recognising new genera among previously congeneric actinopterygians (49), and restriction of *Elonichthys* to just three species (50) rather than its previous 57, will alleviate this issue and mitigate the dominance of superabundant forms. These revisions, however, have the potential to increase unevenness in the other direction, as new taxa may end up as singletons or doubletons. Concurrently, the biostratigraphic significance of actinopterygians in deposits from the Permian of Russia (64–66) may contribute to oversplitting of taxa, echoing problems prevalent in the marine invertebrate fossil record (23).

Major variation in evenness between intervals is highlighted in the different trajectories of coverage-based rarefaction curves (figure S3). Taxonomic and geographic biases are exacerbated by small sample sizes and low coverage, with rarefaction curves crossing multiple times. Higher (more reliable) quorum levels are unobtainable for Palaeozoic actinopterygians due to the high number of singleton taxa (figure S3) controlling Good’s *u* (67). As a result, coverage is generally low (figure S2) and only low quorums—at which evenness signals are more pronounced (24)—can be used. When evenness varies at low levels of sampling, size-based rarefaction can in fact be less biased than coverage-based rarefaction, especially at low levels of coverage (23). Trends between coverage- and size-based rarefaction estimates generally agree (figure S4), although size-based rarefaction estimates higher diversity in some highly uneven intervals (e.g. late Visean; Wuchiapingian and Changhsingian). Small sample sizes (<200 occurrences) also have an effect on the accuracy of coverage estimates using Good’s *u* (23): only four of the sampled Palaeozoic intervals have more than 200 occurrences (late Visean: 266; Serpukhovian: 204; Kasimovian and Gzhelian: 230; Wuchiapingian and Changhsingian: 232). Coverage-based rarefaction curves (figure S3) show these intervals to have among the highest coverage, along with the Bashkirian and Moscovian, highlighting the greater sampling of the Carboniferous than the Permian. Consequently, variation in evenness between intervals is having an overriding effect on sampling-standardised diversity estimates through time, with diversity estimates mostly tracking evenness and reflecting biases in the underlying data (23,68).

## Conclusions and future directions

We present here the first local richness and palaeogeographic trends in Palaeozoic ray-finned fishes. Sampling of the Palaeozoic actinopterygian fossil record is heavily biased towards western Europe (especially the UK) and North America, which translates to a very restricted palaeogeographic spread for most of the Palaeozoic. A suite of compounding problems plagues the actinopterygian fossil record and results in bias towards occurrences of both superabundant and singleton taxa, variation and unevenness in and between sampled intervals, and distortion of relative richness estimates. A combination of flawed taxonomic practices, differential researcher effort, and geographic sampling biases confounds attempts to accurately estimate relative richness between intervals. Meanwhile, sampling is poor for regions other than Europe and North America for all but a few Carboniferous and Permian intervals. The result of this poor sampling is the inability to reach the high levels of coverage that allow statistical methods of sampling standardisation to generate meaningful diversity estimates.

Identifying the underlying issues with Palaeozoic actinopterygian data and the interweaving biases that are impacting the fossil record is crucial, and improving sample sizes and coverage will help to mitigate the sensitivity to evenness (25). Documenting and including existing ‘dark data’ (51,52) in museum collections, as well as focus on new material from under sampled regions, represent key first steps. As a result, size-based rarefaction curves for Palaeozoic intervals will likely not reach asymptote soon (figure S1). More complete sampling of well-known regions (69) may facilitate deduction of accurate local richness patterns (36). This strategy also goes some way towards accounting for the significant spatial structuring of ‘global’ fossil records (17–19,27).

Other recently proposed methods, such as coverage-rarefaction of extrapolated richness estimates (instead of face-value counts) (23), represent prospective avenues of research, both at local and global scales. However, existing global occurrence data for Palaeozoic actinopterygians is as yet inadequate for extrapolation in this way: sample sizes vary widely between intervals, which may result in inaccurate extrapolated richness trends (23,70,71); sample sizes in all intervals are too low for size-based rarefaction curves to asymptote (figure S1), meaning sample size has an overwhelming effect on diversity estimates (23); and abundance distributions are also highly uneven, which biases extrapolators (though to a lesser extent than coverage-based rarefaction; 17).

Overall, the occurrence data recorded in the literature is heavily impacted by sampling and likely results in inaccurate estimated diversity trends at present. Localised diversity estimates for well-sampled regions presents a feasible avenue of research for reconstructing regional diversity. In addition, research efforts to fix problematic taxonomy of ‘waste-basket’ taxa, in hand with a general increase in sampling, open the possibility of estimating diversity in a spatially-standardised framework, so that we can truly begin to understand the origin, rise and establishment of the largest vertebrate clade.

## Acknowledgements

We thank R. Butler, T. Clements, R. Figueroa, M. Friedman, L. Schnetz, R. Close, and G. Lloyd for helpful and insightful discussion during the course of this manuscript. We thank E. Bernard (Natural History Museum, London) and S. Walsh (National Museums of Scotland, Edinburgh) for collections access.

## Funding

This work was funded by a Royal Society Dorothy Hodgkin Research Fellowship (DH160098) and Royal Society Enhancement Award (RGF\EA\180279), both to S. Giles. E. M. Dunne was also supported by a Leverhulme Research Project Grant (RPG-2019-365).

---

**Figure S1.**
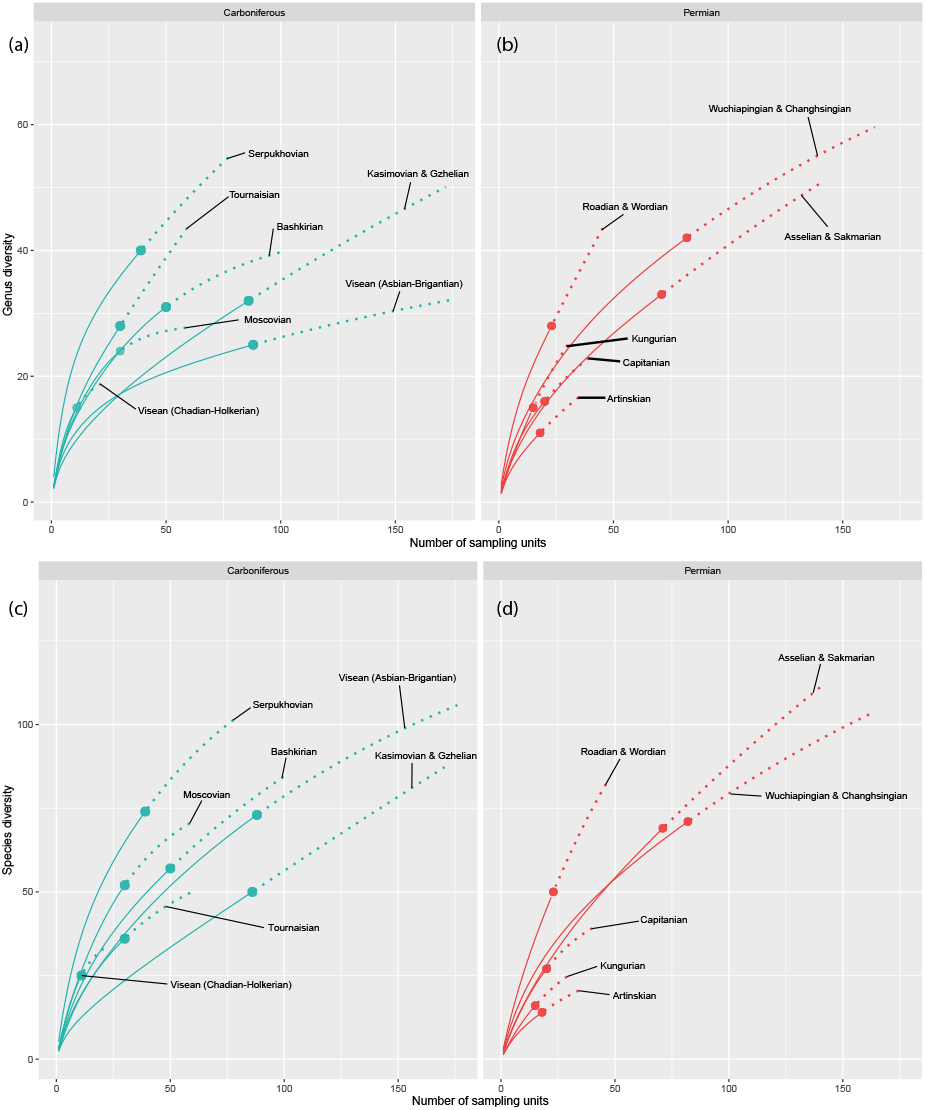
Size-based rarefaction curves for (a) Carboniferous genus-level occurrences, (b) Permian genus-level occurrences, (c) Carboniferous species-level occurrences, (d) Permian genus-level occurrences.

**Figure S2.**
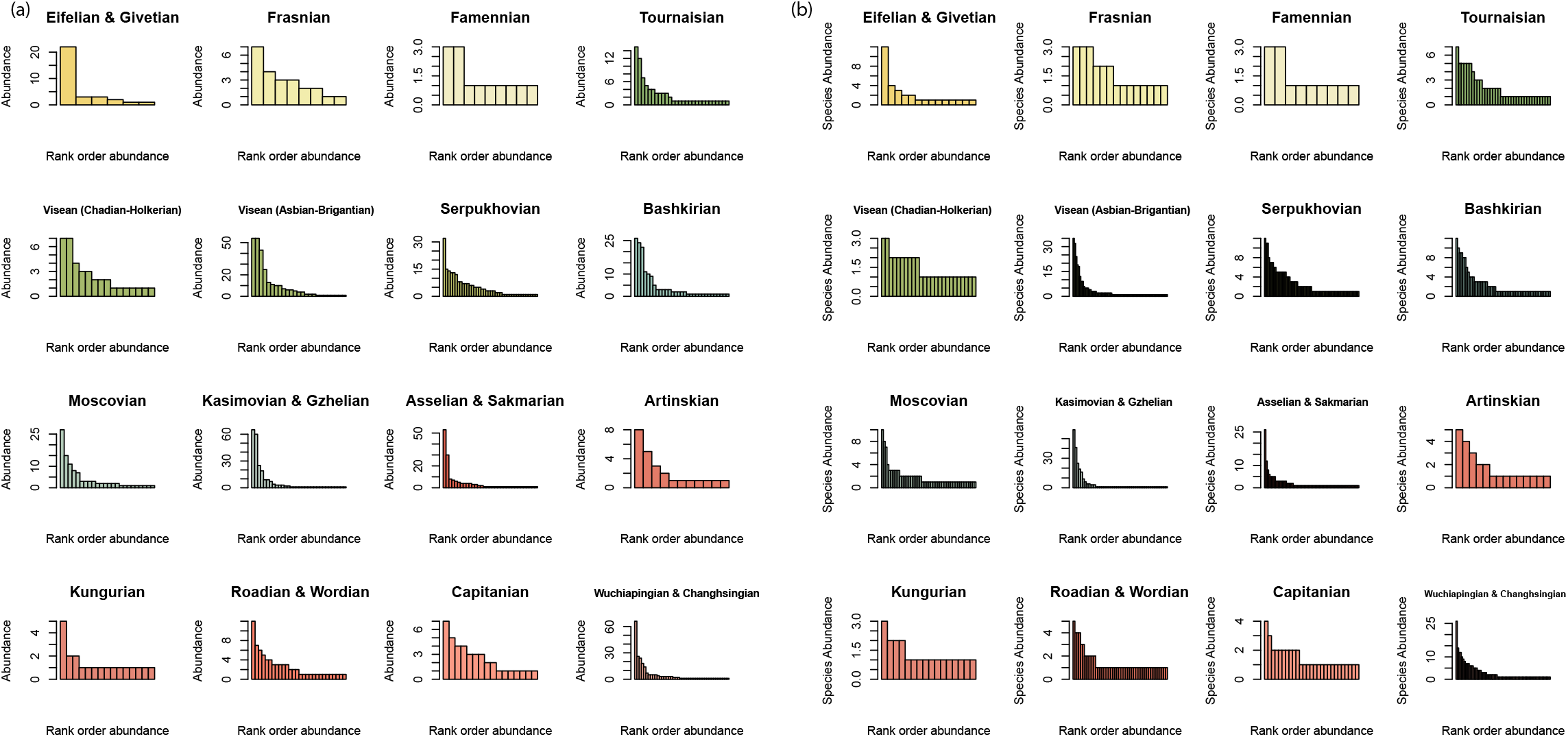
Rank order abundance distributions of the sampled equal-length intervals (coloured correspond with the International Commission on Stratigraphy; ICS) at (a) genus-level occurrences and (b) species-level occurrences.

**Figure S3.**
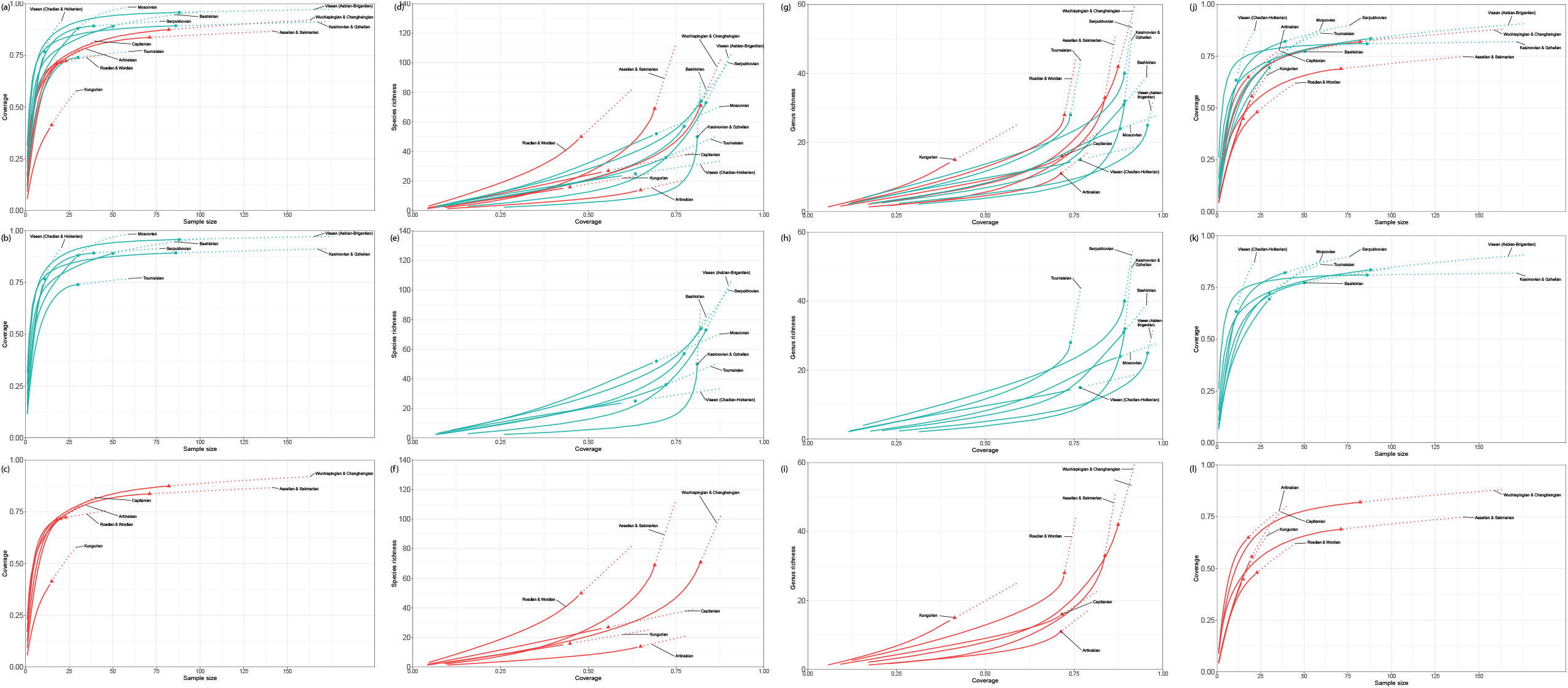
Sample completeness (a-c, k-l) and coverage-based rarefaction curves (d-i) for Carboniferous (b, e, h, k) and Permian (c, f, i, l) intervals. Solid lines are interpolated measures, points are observed data and dotted lines are extrapolated estimates. Carboniferous intervals are in green and Permian intervals are in red according to the ICS colours.

**Figure S4.**
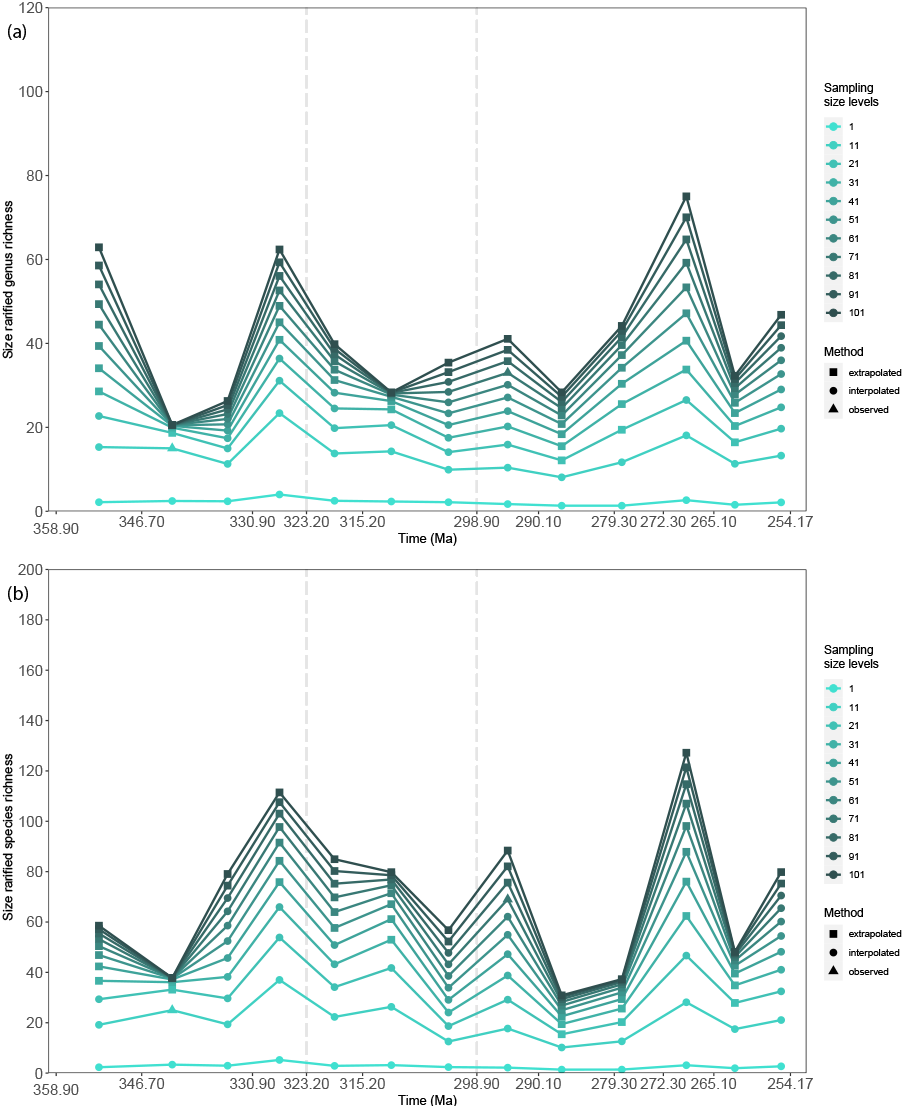
Estimates of Carboniferous and Permian actinopterygian diversity using size-based (classical) rarefaction of (a) occurrences of genera and (b) occurrences of species, showing estimates for different sample sizes.

## Notes

### Competing Interest Statement

The authors have declared no competing interest.

